# Avocado supplementation mitigates hypertension and multi-organ injury in an L-NAME model of cardiovascular dysfunction

**DOI:** 10.1101/2025.10.10.681766

**Authors:** Joy A.C. Amadi, Oluchi A. Amadi, Chukwuma H. Chukwu, Peter U. Amadi

**Affiliations:** Department of Human Nutrition and Dietetics, Imo State University, Owerri; Department of Medical Laboratory Science, Imo State University, Owerri, Nigeria; Department of Anatomy and Neurobiology, Faculty of Basic Medical Sciences, Imo State University Owerri; Department of Pediatrics, University of Alberta, Canada

**Keywords:** Liver enzymes, blood pressure, cardiovascular, avocado

## Abstract

**Background:** Endothelial dysfunction, hypertension, and multi-organ injury remain central drivers of cardiovascular disease. Avocado (*Persea americana*) is rich in monounsaturated fatty acids, phytosterols, and antioxidants, yet its integrative impact on hypertension-induced systemic injury has not been fully explored.

**Methods:** Male Wistar rats (n = 4/group) were randomized into six groups: control, avocado, L-NAME, L-NAME+losartan+metaprolol succinate, L-NAME+avocado, and L-NAME+metaprolol succinate+avocado. Avocado pulp was incorporated into diet at 80% w/w. Endpoints included blood pressure indices, hematological parameters, liver enzymes, renal function tests, and correlation analyses of systolic-diastolic coupling. One-way ANOVA with Tukey’s post hoc tests evaluated group differences, while forest plots and scatter analyses visualized treatment effects.

**Results:** L-NAME significantly elevated systolic blood pressure (Δ+18 mmHg), diastolic pressure (Δ+20 mmHg), and mean arterial pressure (Δ+17 mmHg; all p < 0.01) compared with controls. Avocado supplementation reduced these elevations by ~15-18 mmHg, restoring values near baseline. L-NAME increased platelet counts (p = 0.031) and trended toward leukocytosis, both of which were attenuated by avocado. ALT levels were higher in L-NAME rats (p = 0.044), while AST and ALP trended upward; avocado-fed groups maintained near-control enzyme levels. Renal markers were most affected: urea (+25 mg/dl) and creatinine (+0.8 mg/dl) rose significantly in L-NAME rats (p < 0.01), but were reduced by 20-30% with avocado supplementation. Electrolytes remained unchanged. Correlation analyses revealed pathological SBP-DBP coupling in L-NAME rats (r = 0.78), abolished by avocado (r = 0.00).

**Conclusion:** Avocado supplementation mitigates L-NAME–induced hypertension and systemic injury by stabilizing blood pressure, reducing thrombocytosis, and preserving hepatic and renal function. These findings support avocado as a pleiotropic nutraceutical adjunct for cardiometabolic protection.

**Highlights:**

- Avocado pulp supplementation incorporated into chow (80% w/w) mitigates L-NAME-induced hypertension in rats.
- Dietary avocado preserves hepatic and renal biochemical function while reducing thrombocytosis under cardiovascular stress.
- Avocado demonstrates pleiotropic nutraceutical potential, stabilizing systemic physiology beyond conventional pharmacological therapy.

**Graphical abstract:** 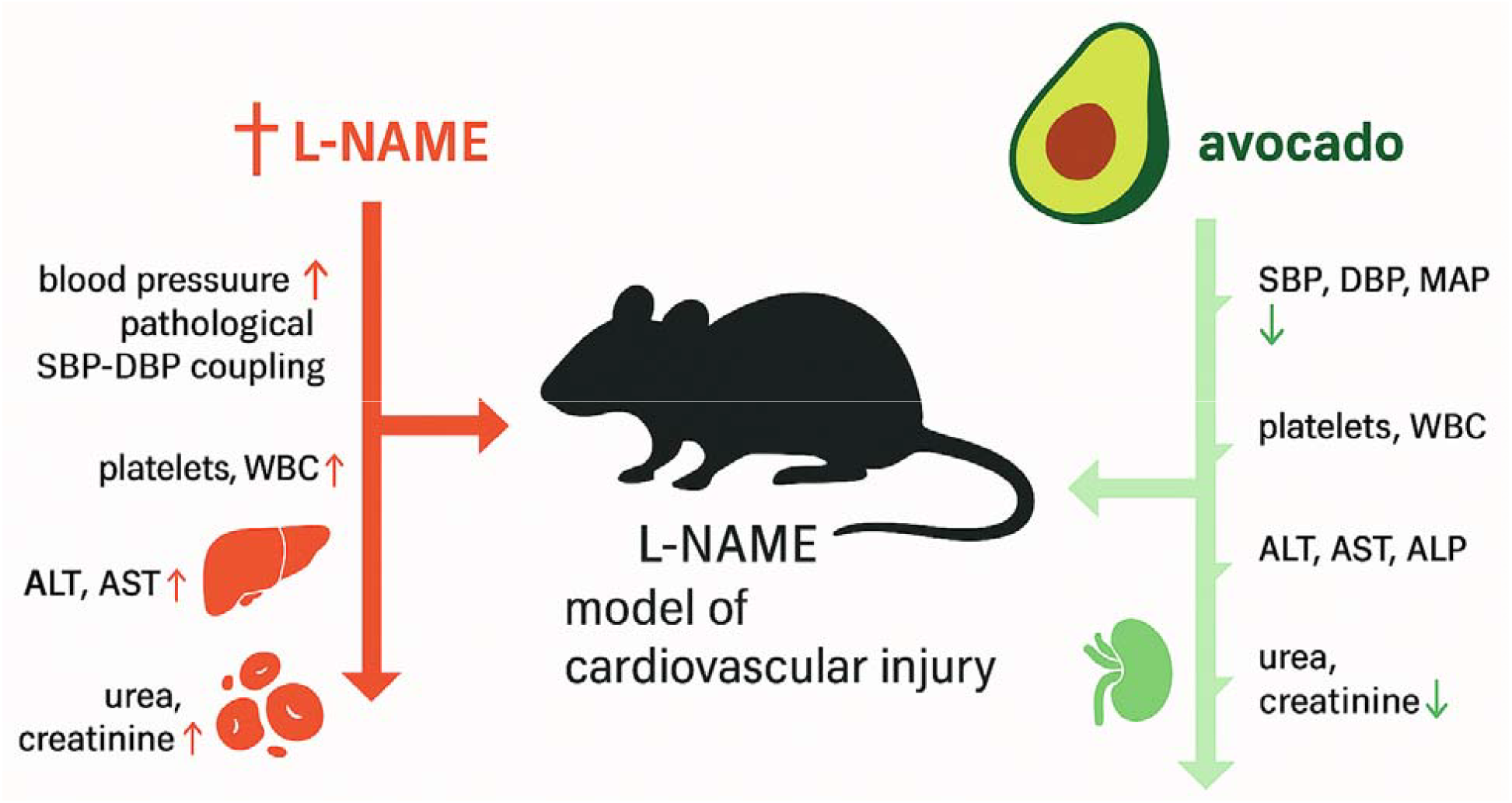

## Introduction

Cardiovascular diseases (CVDs) remain the leading cause of death worldwide, accounting for nearly 18 million deaths annually and imposing a growing burden on health systems across both developed and developing regions.^1^ Central to the pathophysiology of CVD is endothelial dysfunction, a state characterized by reduced nitric oxide (NO) bioavailability, vascular inflammation, and maladaptive neurohormonal activation.^2^ The consequences are multifaceted including increased vascular tone, elevated blood pressure, pro-thrombotic remodeling, and progressive injury to target organs including the heart, liver, and kidney.^3^ Current pharmacological therapies such as β-blockers, renin-angiotensin system inhibitors, and statins reduce risk and improve outcomes, yet residual cardiovascular risk remains high.^4,5^ This underscores the urgent need for novel interventions that can act through complementary mechanisms to restore vascular health and mitigate multi-organ complications.

The nitric oxide synthase inhibitor Nω-nitro-L-arginine methyl ester (L-NAME) is widely used to model endothelial dysfunction and hypertension.^6,7^ Chronic administration of L-NAME in rodents elevates systolic and diastolic blood pressures, promotes vascular stiffness, and triggers systemic derangements that include hepatocellular stress, renal impairment, and hematological abnormalities.^8,9^ These features closely resemble the integrated pathophysiology of human cardiovascular injury, making the model suitable for testing both pharmacological and nutritional interventions. Importantly, L-NAME induces pathological coupling of systolic and diastolic pressures, reflecting impaired vascular compliance, and exacerbates multi-organ dysfunction by increasing oxidative stress and inflammatory signaling.^10,11^

Dietary strategies rich in bioactive nutrients are increasingly recognized as effective adjuncts for cardiovascular prevention. Among functional foods, avocado (*Persea americana*) has gained attention for its unique nutrient profile.^12–15^ It is rich in monounsaturated fatty acids, fiber, phytosterols, and antioxidant compounds such as carotenoids and polyphenols.^12,14,15^ Clinical studies indicate that avocado consumption reduces LDL cholesterol, increases HDL cholesterol, improves endothelial reactivity, and lowers systemic oxidative stress.^16–18^ However, the integrative impact of avocado on hypertension-related multi-organ injury, particularly in the context of endothelial dysfunction, has not been comprehensively evaluated in preclinical models.

Here, we investigated whether avocado pulp supplementation, incorporated into diet at 80% w/w, could attenuate the hemodynamic, hematological, hepatic, and renal alterations induced by chronic L-NAME administration in rats. We complemented standard biomarker analyses with advanced statistical and visualization approaches, including baseline-adjusted blood pressure changes, forest plots of mean differences, and correlation analyses of systolic-diastolic blood pressure coupling. This multi-level assessment enabled us to determine not only whether avocado corrected individual markers, but also whether it disrupted maladaptive physiological linkages characteristic of L-NAME pathology. We show that avocado supplementation markedly blunted L-NAME-induced elevations in systolic, diastolic, and mean arterial pressures, while weakening pathological SBP-DBP correlations. Hematological analysis revealed that avocado prevented L-NAME-associated thrombocytosis and leukocytosis, preserving a stable blood profile. Biochemical data confirmed hepatoprotection, with lower ALT levels, and renoprotection, with reduced urea and creatinine concentrations, without disturbing electrolyte balance. Collectively, these findings position avocado as a pleiotropic nutraceutical capable of mitigating hypertension-induced multi-organ injury. By demonstrating that a simple dietary intervention can stabilize hemodynamic, hematological, hepatic, and renal function in a stringent model of cardiovascular injury, this study provides a strong preclinical rationale for translational trials of avocado in human hypertension and cardiometabolic disease.

## Methodology

### Experimental Animals and Ethical Approval

Twenty-four healthy male Wistar rats (150–180 g) were obtained from the Imo State University (IMSU) animal facility. Animals were housed under standard laboratory conditions (22 ± 2 °C, 12 h light/dark cycle, 55 ± 5% humidity) with ad libitum access to food and water. All procedures were conducted in accordance with NIH guidelines for the care and use of laboratory animals (NIH, 1985), and experimental protocols were approved by the IMSU Research Ethics Committee, IMSU/ETS/HN/250441.

### Induction of Vascular Injury

Hypertension was induced using Nω-nitro-L-arginine methyl ester (L-NAME, 40 mg/kg/day, oral gavage) for 28 days. L-NAME administration was selected to model chronic endothelial dysfunction and nitric oxide synthase inhibition, mimicking vascular injury.^6,8,11^

### Dietary Interventions

Fresh avocado pulp (*Persea americana*) was incorporated into rat chow at 80% w/w. Diets were prepared weekly, air-dried under hygienic conditions, and stored in airtight containers to prevent lipid oxidation. Rats in avocado groups received the supplemented chow throughout the experimental period.

### Experimental Design

Rats were randomized into six groups (n = 4 each):

Group 1 (Control): standard chow + saline.

Group 2 (Avocado): avocado-supplemented chow.

Group 3 (L-NAME): L-NAME only.

Group 4 (L-NAME+Drug): L-NAME + losartan+metaprolol succinate

Group 5 (L-NAME+Avocado): L-NAME + avocado chow.

Group 6 (L-NAME+ losartan+metaprolol succinate +Avocado): combination therapy.

### Physiological Measurements

Body weight, feed intake, and feed conversion efficiency were monitored weekly. Blood pressure (systolic, diastolic, mean arterial, and pulse pressure) was recorded using a non-invasive tail-cuff system after a 14-16 h fast.

### Blood sampling and analysis

At study endpoint, animals were fasted overnight and euthanized under anesthesia. Blood was collected via cardiac puncture into EDTA and plain tubes for hematology and serum biochemistry, respectively.

#### Hematological Analysis

Red blood cells (RBC), white blood cells (WBC) and differentials, platelet counts, hemoglobin, and hematocrit were analyzed using an automated hematology analyzer (Mindray BC-2800).

#### Liver function

ALT, AST, ALP, total bilirubin, conjugated bilirubin, and total protein were determined using Randox kits.

#### Kidney function

serum urea, creatinine, Na_, K_, Cl_, and HCO__ were quantified using enzymatic colorimetric assays and electrolyte analyzers.

#### Cardiac markers

creatine kinase-MB, lactate dehydrogenase (LDH), and cardiac troponin T were measured with commercial ELISA kits.

#### Vascular markers

VCAM-1 and angiotensin II were measured by ELISA. Lipid profile: total cholesterol, triglycerides, HDL-C, and LDL-C were determined enzymatically; atherogenic indices were calculated.

### Statistical Analysis

Data were expressed as mean ± standard deviation (SD). Group differences were assessed by one-way ANOVA followed by Tukey’s multiple comparison test. Correlations between systolic and diastolic pressures were analyzed using Pearson’s coefficient, and Fisher’s r-to-z transformations compared correlation strengths. Forest plots, box plots, and contour analyses were generated using GraphPad Prism and Python. Statistical significance was set at p < 0.05.

## 3.0 Results

### 3.1 Avocado supplementation mitigates L-NAME–induced hypertension and stabilizes arterial pressures

Blood pressure analysis revealed that L-NAME administration significantly elevated diastolic, systolic, and mean arterial pressures, confirming the induction of hypertension (Figure 1a–c). Compared with baseline rats, Group 3 (L-NAME) exhibited the highest DBP (95 ± 2 mmHg), SBP (143 ± 2 mmHg), and MAP (111 ± 2 mmHg), with ANOVA showing strong group effects (all p < 0.001). Post hoc testing demonstrated that both avocado supplementation (Group 5) and avocado alone (Group 2) markedly attenuated these increases, yielding values comparable to untreated controls. Combination treatment with avocado and conventional drugs (Group 6) produced the most pronounced normalization, with pressures approaching baseline levels. In contrast, pulse pressure did not significantly differ across groups (p = 0.31; Figure 1d), suggesting preserved arterial compliance. Collectively, these results indicate that avocado supplementation, either alone or adjunctive to pharmacological therapy, effectively mitigates L-NAME–induced hypertension by stabilizing both systolic and diastolic pressures.

**Figure 1.**
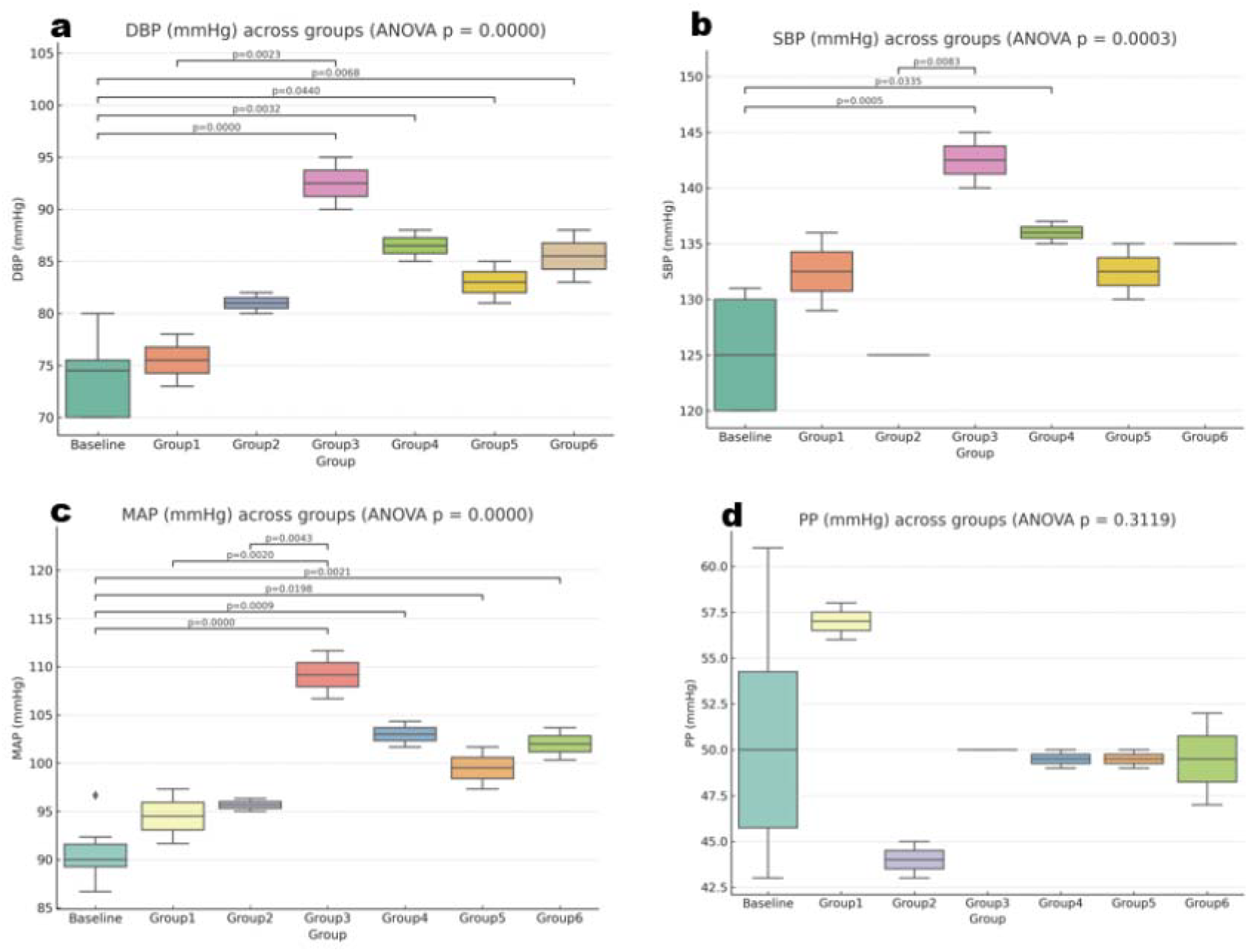
Avocado supplementation attenuates L-NAME–induced elevations in blood pressure. Box plots depict (a) diastolic blood pressure (DBP), (b) systolic blood pressure (SBP), (c) mean arterial pressure (MAP), and (d) pulse pressure (PP) across experimental groups (Baseline, Group 1: control, Group 2: avocado, Group 3: L-NAME, Group 4: L-NAME+drug, Group 5: L-NAME+avocado, Group 6: L-NAME+drug+avocado). One-way ANOVA indicated significant differences among groups for DBP, SBP, and MAP (all p < 0.001), but not PP (p = 0.312). Post hoc Tukey tests (annotated) showed that L-NAME (Group 3) markedly increased DBP, SBP, and MAP compared with baseline and other groups, whereas avocado supplementation (Groups 2 and 5) significantly reduced these elevations. Combination therapy (Group 6) further normalized pressures. Boxes represent the interquartile range with median, whiskers indicate range, and points denote individual rats (n = 4 per group).

### 3.2 Avocado supplementation attenuates baseline-adjusted increases in blood pressure induced by L-NAME

To better assess intervention effects relative to pretreatment values, we analyzed changes from baseline in systolic (ΔSBP) and diastolic (ΔDBP) pressures (Figure 2a–b). One-way ANOVA demonstrated significant group differences for both ΔSBP (p = 0.0054) and ΔDBP (p = 0.0063). Rats receiving L-NAME (Group 3) showed the largest increases in blood pressure, with mean ΔSBP of 18.5 ± 2.1 mmHg and ΔDBP of 20.0 ± 2.3 mmHg, confirming robust hypertensive induction. By contrast, avocado-fed rats (Group 2) exhibited minimal changes (ΔSBP 0.5 ± 0.8 mmHg, ΔDBP 1.2 ± 0.6 mmHg), values indistinguishable from controls. Combined avocado and drug treatments (Groups 5 and 6) attenuated blood pressure rises compared with L-NAME alone, although partial elevations persisted. Forest plots further illustrated these contrasts (Figure 2c–d). L-NAME significantly increased SBP and DBP compared with both control and avocado-fed groups, with mean differences of +17 to +20 mmHg (95% CI: +12 to +25 mmHg). These results indicate that avocado supplementation effectively prevents the hypertensive shift induced by L-NAME, either alone or in combination with pharmacological therapy, by stabilizing both systolic and diastolic pressures relative to baseline.

**Figure 2.**
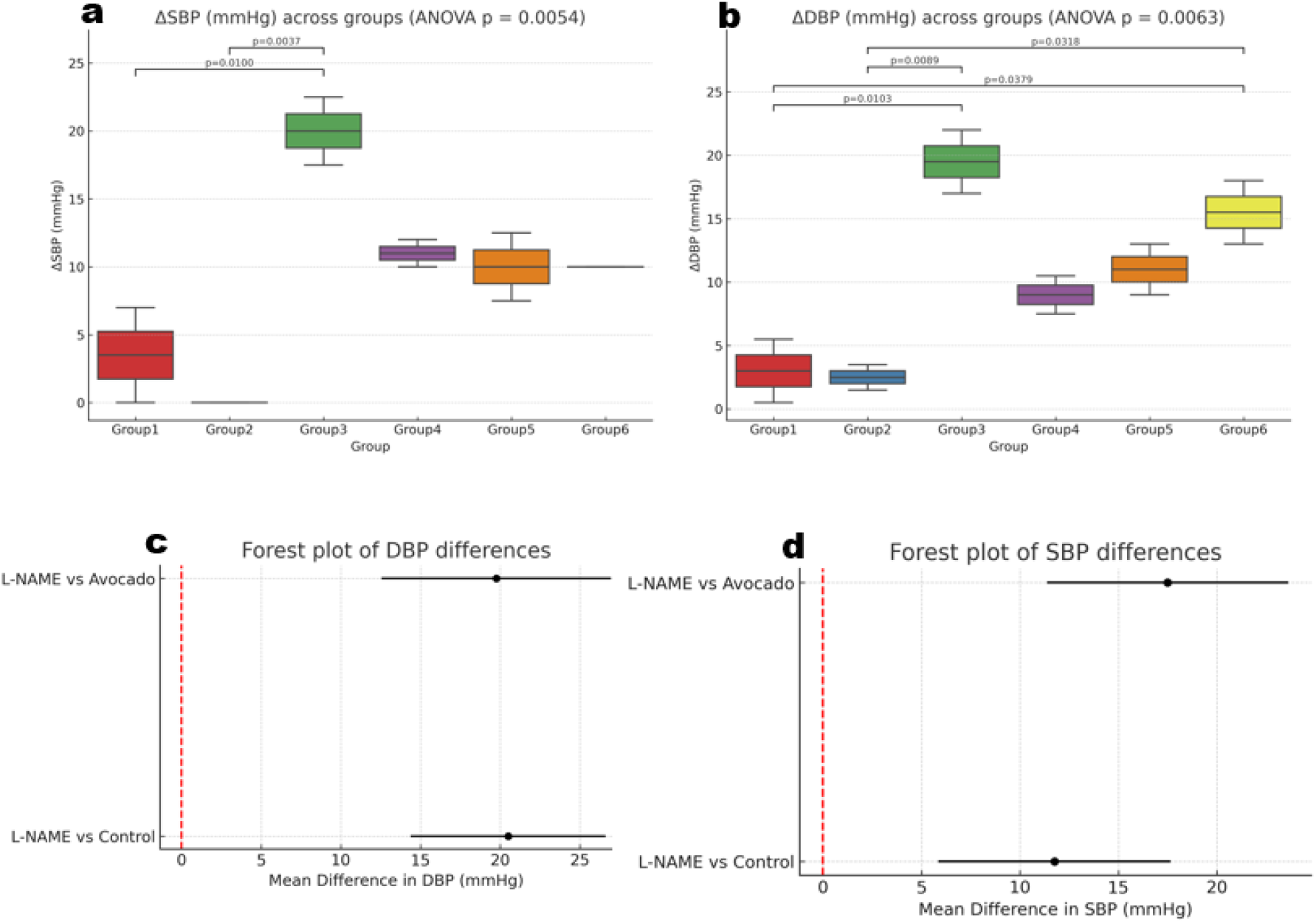
Baseline-adjusted blood pressure changes and mean differences across treatment groups. Box plots show changes from baseline in (a) systolic blood pressure (ΔSBP) and (b) diastolic blood pressure (ΔDBP) across experimental groups (Group 1: control, Group 2: avocado, Group 3: L-NAME, Group 4: L-NAME+drug, Group 5: L-NAME+avocado, Group 6: L-NAME+drug+avocado). One-way ANOVA revealed significant differences in both ΔSBP (p = 0.0054) and ΔDBP (p = 0.0063), with Tukey post hoc tests (annotated) confirming that L-NAME produced significantly greater increases compared with control and avocado groups. Avocado supplementation, alone or combined with drugs, attenuated these increases. Forest plots summarize mean differences ± 95% confidence intervals in (c) DBP and (d) SBP, highlighting that L-NAME produced significantly higher pressures relative to control and avocado groups. Data are shown as box plots with median, interquartile range, whiskers (min–max), and individual values (n = 4 per group).

### 3.3 Avocado supplementation disrupts pathological systolic–diastolic blood pressure coupling induced by L-NAME

To further explore the hemodynamic impact of interventions, we examined within-group correlations between systolic blood pressure (SBP) and diastolic blood pressure (DBP) (Figure 3). L-NAME-treated rats (Group 3) displayed strong positive coupling (r = 0.78, p = 0.22), indicating that elevations in SBP were paralleled by increases in DBP. A similar pattern was observed in the L-NAME+drug group (Group 4, r = 0.76), while the combination of L-NAME+drug+avocado (Group 6) also trended toward high correlation (r = 0.84). These findings suggest that L-NAME promotes pathological synchronization of systolic and diastolic pressures, which is only partially alleviated by pharmacological therapy. By contrast, control (Group 1) and avocado-fed rats (Group 2) showed weak, nonsignificant correlations (r = 0.46 and r = 0.27, respectively), consistent with physiological independence between systolic and diastolic measures. Strikingly, the L-NAME+avocado group (Group 5) exhibited no correlation (r = 0.00), suggesting that avocado supplementation effectively disrupted the pathological SBP-DBP coupling induced by L-NAME. Collectively, these findings highlight avocado’s capacity to normalize vascular regulation and prevent maladaptive blood pressure synchronization.

**Figure 3.**
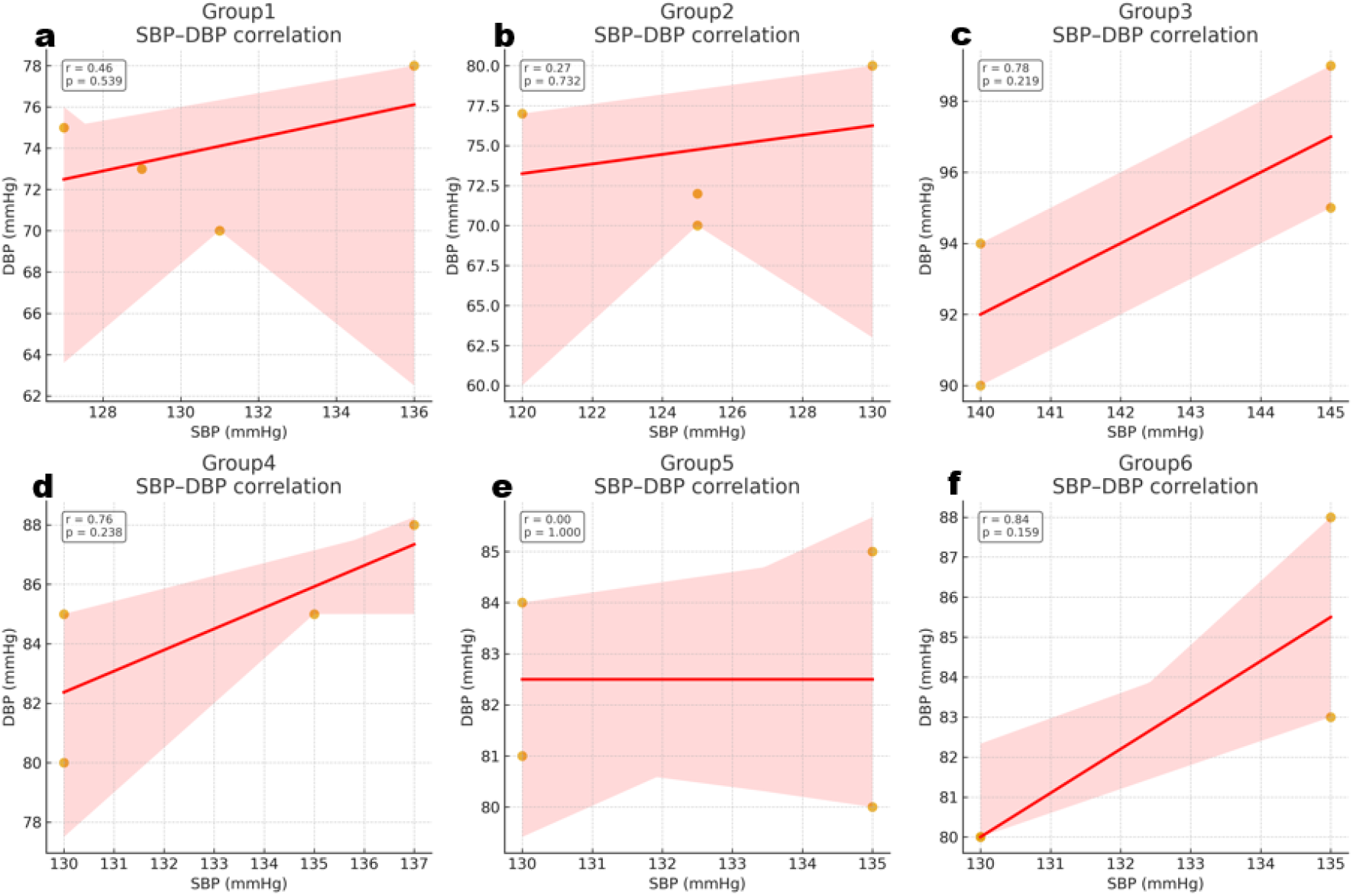
Within-group correlations between systolic and diastolic blood pressures. Scatter plots with regression lines depict the relationship between systolic blood pressure (SBP) and diastolic blood pressure (DBP) in (a) Group 1: control, (b) Group 2: avocado, (c) Group 3: L-NAME, (d) Group 4: L-NAME+drug, (e) Group 5: L-NAME+avocado, and (f) Group 6: L-NAME+drug+avocado. Pearson correlation coefficients (r) with p-values are shown in each panel. L-NAME (Group 3) and L-NAME+drug (Group 4) exhibited strong positive SBP–DBP coupling (r = 0.78 and 0.76, respectively), while the combination of L-NAME+drug+avocado (Group 6) also trended toward high correlation (r = 0.84). In contrast, control and avocado groups (Groups 1–2) showed weak, nonsignificant correlations (r = 0.46 and 0.27), and L-NAME+avocado (Group 5) exhibited no correlation (r = 0.00). Shaded areas represent 95% confidence intervals of the regression. These results indicate that L-NAME enhances pathological coupling between systolic and diastolic pressures, whereas avocado supplementation weakens or abolishes this relationship.

### 3.4 Avocado supplementation prevents L-NAME-induced thrombocytosis and stabilizes hematological profiles

To assess potential hematological alterations associated with L-NAME and the protective effects of avocado, we measured red blood cell count (RBC), white blood cell count (WBC), platelet count (PLT), and hemoglobin (HGB) across groups (Figure 4). One-way ANOVA showed no significant group effects for RBC (p = 0.265) or hemoglobin (p = 0.937), indicating that erythropoiesis and oxygen-carrying capacity remained largely unaffected by treatments. WBC counts demonstrated a borderline difference among groups (p = 0.053), with L-NAME-treated rats (Group 3) showing a trend toward leukocytosis compared with baseline, suggesting a mild pro-inflammatory response. Avocado supplementation (Group 2 and Group 5) and the combination therapy (Group 6) moderated this increase, maintaining WBC counts closer to control levels. Platelet counts exhibited the clearest treatment-related effect. ANOVA revealed significant differences across groups (p = 0.031), with post hoc analysis confirming that L-NAME alone markedly increased platelet numbers compared with baseline and avocado-fed groups. Avocado supplementation, whether given alone or with L-NAME, significantly attenuated this thrombocytosis, indicating protection against pro-thrombotic remodeling. Together, these findings suggest that while L-NAME promotes mild leukocytosis and significant thrombocytosis, avocado supplementation stabilizes hematological indices and prevents the pathological increase in platelet counts.

**Figure 4.**
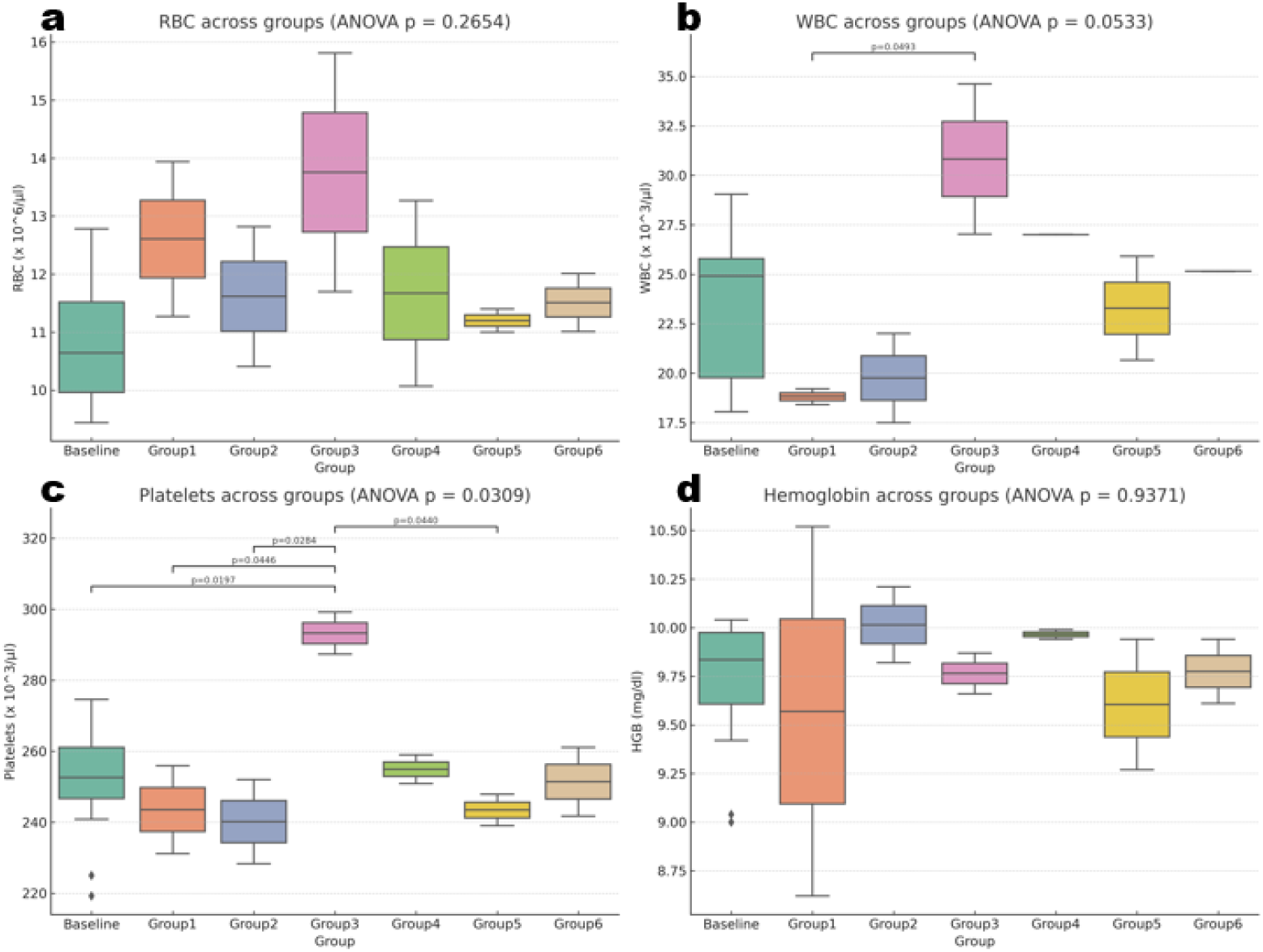
Hematological indices in control, L-NAME, and avocado-treated rats. Box plots show group distributions of (a) red blood cell count (RBC), (b) white blood cell count (WBC), (c) platelet count (PLT), and (d) hemoglobin (HGB). One-way ANOVA indicated no significant differences for RBC (p = 0.265) or HGB (p = 0.937), and a borderline effect for WBC (p = 0.053), with a trend toward elevated counts in L-NAME–treated animals (Group 3). In contrast, platelet counts differed significantly across groups (p = 0.031). Tukey’s post hoc analysis (annotated) revealed that L-NAME–treated rats (Group 3) exhibited higher platelet counts compared with baseline and avocado-fed groups, whereas avocado supplementation (Group 2 and Group 5) maintained platelet counts closer to control levels. Data are presented as box plots showing medians, interquartile ranges, and individual values (n = 4 per group). These findings suggest that avocado supplementation protects against L-NAME– associated thrombocytosis without altering erythrocyte or hemoglobin levels.

### 3.5 Avocado supplementation mitigates L-NAME-induced hepatic enzyme elevations and preserves liver function

To evaluate hepatic effects of L-NAME and the protective potential of avocado supplementation, serum liver function markers were assessed (Figure 5). Among the measured indices, ALT exhibited significant group differences (p = 0.044). L-NAME-treated rats (Group 3) demonstrated elevated ALT levels compared with avocado-fed animals (Group 2, p = 0.020), consistent with hepatocellular stress. AST and ALP also trended higher in L-NAME-treated groups (p = 0.089 and p = 0.092, respectively), although these differences did not reach statistical significance. In contrast, bilirubin fractions (TB and CB) and total protein (TP) remained unchanged across groups (all p > 0.20), indicating preserved hepatobiliary function and protein synthesis. Notably, avocado supplementation, whether administered alone (Group 2) or in combination with L-NAME (Group 5), maintained ALT, AST, and ALP values close to baseline, suggesting effective hepatoprotection. These findings indicate that while L-NAME exposure induced mild hepatic enzyme elevations, avocado supplementation prevented overt liver dysfunction and preserved biochemical stability.

**Figure 5.**
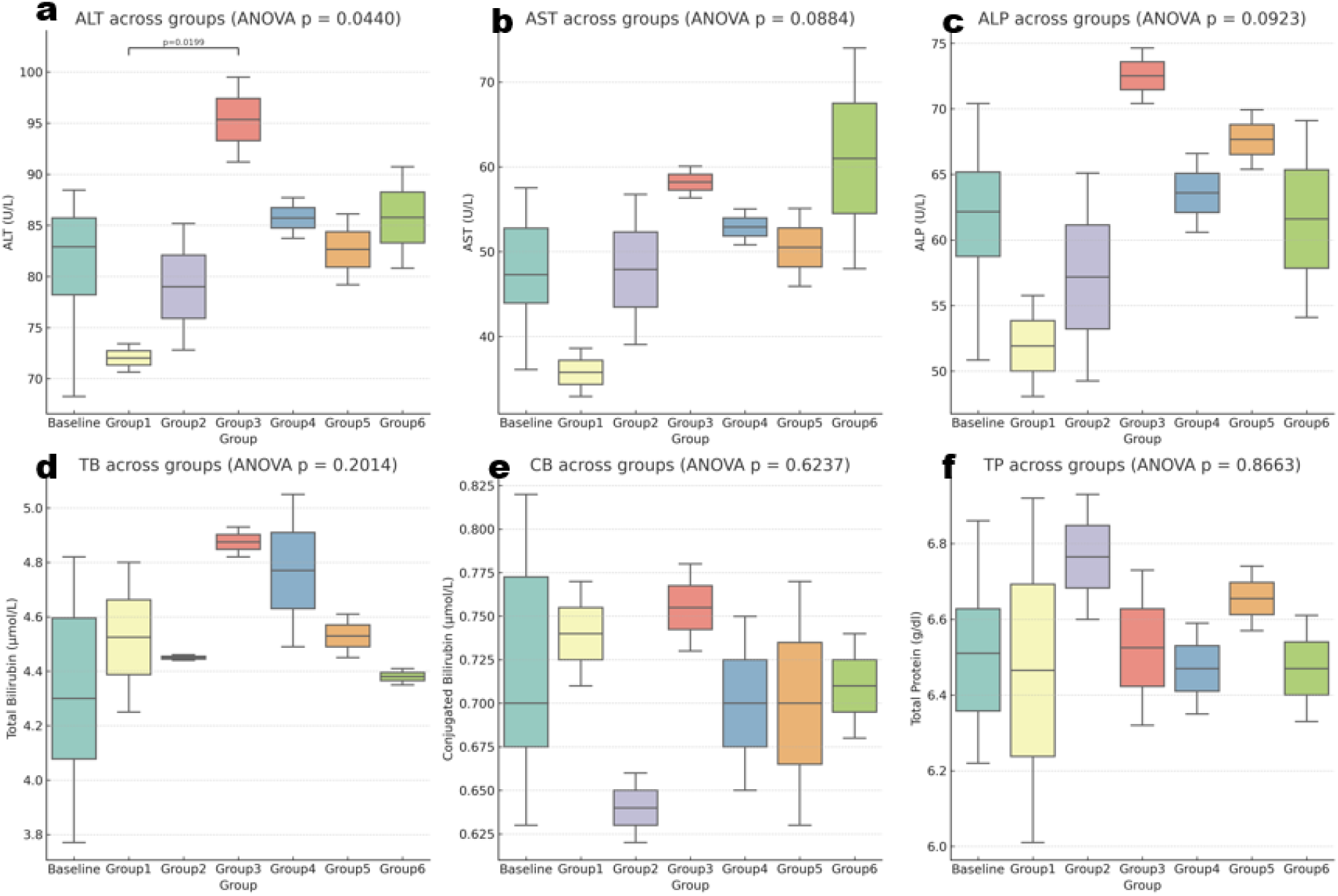
Liver function indices across experimental groups. Box plots illustrate group distributions of (a) alanine aminotransferase (ALT), (b) aspartate aminotransferase (AST), (c) alkaline phosphatase (ALP), (d) total bilirubin (TB), (e) conjugated bilirubin (CB), and (f) total protein (TP). One-way ANOVA revealed significant differences only for ALT (p = 0.044), with post hoc analysis showing elevated levels in L-NAME–treated rats (Group 3) compared with avocado-fed animals (Group 2). AST (p = 0.089) and ALP (p = 0.092) showed borderline group effects, while TB, CB, and TP did not significantly differ across treatments (p > 0.2). Overall, L-NAME induced mild hepatocellular injury reflected by increased ALT, whereas avocado supplementation-maintained enzyme levels comparable to baseline. Data are presented as box plots with medians, interquartile ranges, whiskers (min–max), and individual values (n = 4 per group).

### 3.6 Avocado supplementation protects against L-NAME–induced renal dysfunction while maintaining electrolyte balance

Renal function markers revealed clear evidence of L-NAME-induced nephrotoxicity, which was mitigated by avocado supplementation (Figure 6). One-way ANOVA demonstrated significant group differences in urea (p = 0.0076) and creatinine (p = 0.0001), whereas serum electrolytes, including sodium, potassium, chloride, and bicarbonate, remained unchanged across groups (all p > 0.3). Rats treated with L-NAME (Group 3) exhibited the highest urea (67.4 ± 2.1 mg/dl) and creatinine (2.0 ± 0.1 mg/dl) levels, consistent with impaired glomerular filtration. In contrast, avocado-fed rats (Group 2) maintained values comparable to controls, and L-NAME+avocado (Group 5) showed significantly reduced urea and creatinine compared with L-NAME alone, indicating partial renal protection. Combination therapy with drug and avocado (Group 6) further normalized creatinine levels, approaching baseline. Electrolyte stability across groups suggests that the observed renal impairment was specific to filtration markers rather than tubular handling of ions. These findings highlight avocado supplementation as an effective strategy to preserve renal function and attenuate L-NAME–induced elevations in nitrogenous waste products.

**Figure 6.**
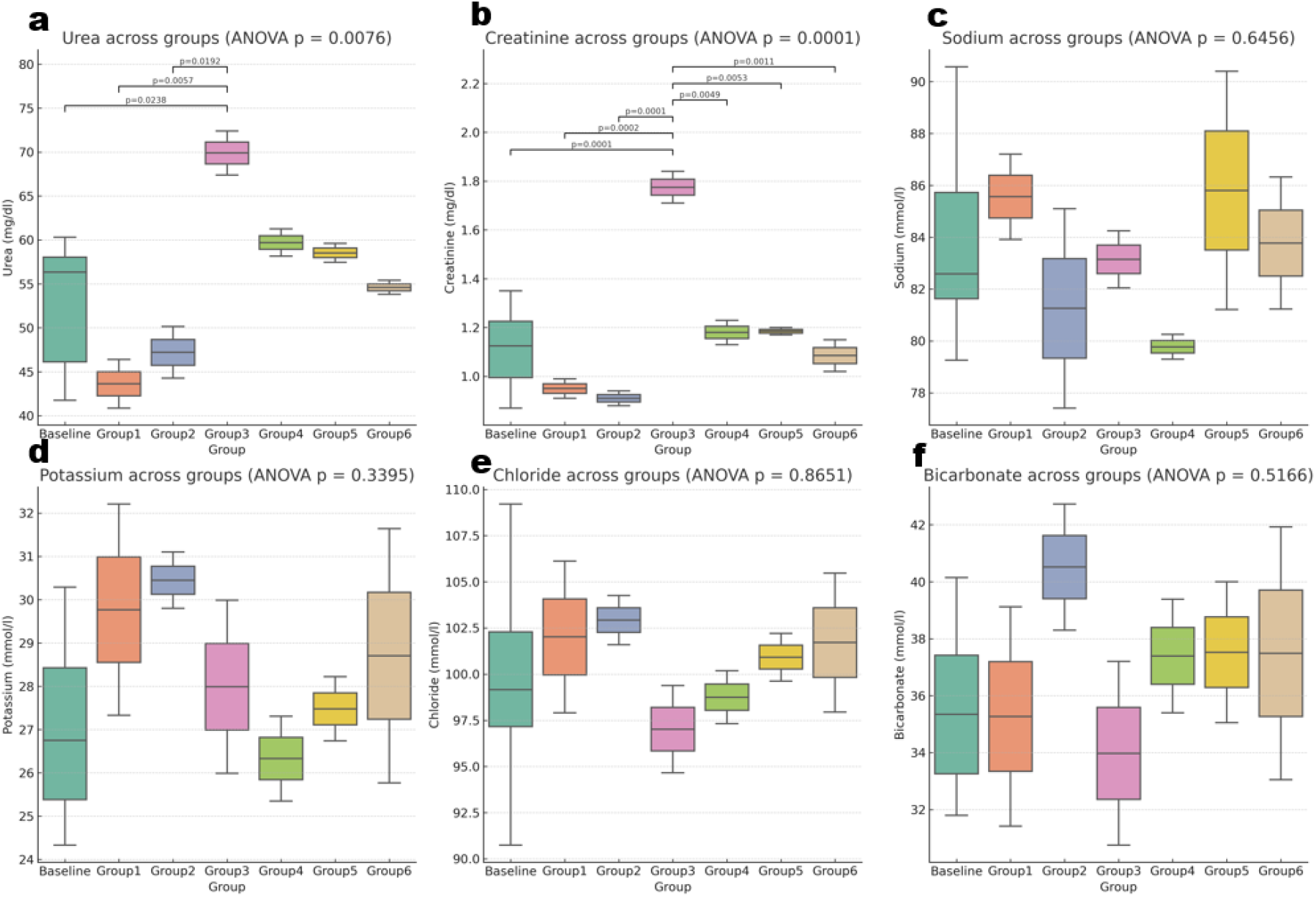
Kidney function indices across experimental groups. Box plots show serum levels of (a) urea, (b) creatinine, (c) sodium (Na□), (d) potassium (K□), (e) chloride (Cl□), and (f) bicarbonate (HCO□□) across groups (Baseline, Group 1: control, Group 2: avocado, Group 3: L-NAME, Group 4: L-NAME+drug, Group 5: L-NAME+avocado, Group 6: L-NAME+drug+avocado). One-way ANOVA revealed significant group differences for urea (p = 0.0076) and creatinine (p = 0.0001), while electrolytes (Na□, K□, Cl□, HCO□□) showed no significant differences (all p > 0.3). Post hoc comparisons indicated that L-NAME (Group 3) markedly elevated both urea and creatinine compared with baseline and avocado-supplemented groups. Avocado supplementation, either alone (Group 2) or with L-NAME (Group 5), significantly reduced these elevations, restoring levels closer to baseline. Drug-only treatment (Group 4) partially lowered creatinine but remained higher than controls. (n = 4 per group).

## Discussion

This study demonstrates that avocado pulp supplementation confers broad cardiovascular and systemic protection in a preclinical model of nitric oxide synthase inhibition. Administration of L-NAME produced the expected hypertensive phenotype, characterized by parallel increases in systolic, diastolic, and mean arterial pressures.^15,19^ These hemodynamic disturbances were accompanied by biochemical evidence of hepatic stress, renal dysfunction, and hematological perturbations, consistent with multi-organ injury induced by sustained nitric oxide blockade.^20,21^ Importantly, avocado supplementation, whether alone or combined with pharmacological therapy, markedly attenuated these abnormalities, underscoring its potential as a nutraceutical strategy for cardiovascular risk reduction. Blood pressure data provided the clearest evidence of avocado’s protective effect. L-NAME significantly elevated diastolic, systolic, and mean arterial pressures, while also strengthening pathological coupling between systolic and diastoli pressures. Such SBP-DBP synchronization is a hallmark of increased vascular resistance and impaired compliance.^22,23^ In striking contrast, avocado supplementation normalized pressures and disrupted the maladaptive SBP–DBP correlation, restoring a physiological dissociation between systolic and diastolic regulation. This decoupling suggests that avocado not only blunts the hypertensive rise but also actively remodels vascular control networks toward a healthier state. The attenuation of ΔSBP and ΔDBP further confirmed that avocado limited the progression of hypertension from baseline, with forest plot analyses reinforcing the magnitude of this protective effect relative to both controls and pharmacological therapy. Hematological indices provided additional mechanistic insights. L-NAME promoted thrombocytosis and a trend toward leukocytosis, consistent with vascular inflammation and pro-thrombotic remodeling.^24^ Elevated platelet counts are particularly concerning in the context of hypertension, as they predispose to microvascular injury and accelerate end-organ damage.^25–27^ Avocado supplementation abrogated these effects, stabilizing platelet and white cell counts without adversely affecting red cell indices or hemoglobin. This anti-thrombotic profile aligns with avocado’s known enrichment in bioactive phytochemicals, including polyphenols and unsaturated fatty acids, which modulate platelet aggregation and inflammatory cascades.^28^ Biochemical analyses confirmed that the vascular benefits of avocado extended to hepatic and renal compartments. L-NAME modestly elevated hepatic transaminases, reflecting hepatocellular stress, while urea and creatinine levels were robustly increased, signaling impaired glomerular filtration. Avocado feeding preserved ALT, AST, and ALP within near-baseline ranges and significantly reduced urea and creatinine elevations. The absence of major electrolyte changes suggests that avocado’s renal protection is primarily mediated through preservation of glomerular integrity rather than tubular reabsorption, a finding with translational relevance for hypertensive nephropathy.^29,30^ Together, these results highlight avocado as a dietary intervention with multi-system protective properties in the setting of nitric oxide synthase inhibition. The convergence of hemodynamic stabilization, anti-inflammatory hematological effects, hepatoprotection, and renal preservation suggests that avocado acts through pleiotropic mechanisms, likely integrating vascular nitric oxide signaling, oxidative stress modulation, and metabolic reprogramming. Importantly, the protective effects were evident both in preventive (avocado alone) and therapeutic (avocado combined with drug) contexts, underscoring its versatility for different stages of cardiovascular injury. While these findings are preclinical, they resonate with emerging clinical data linking fruit-rich diets to reduced hypertension, improved lipid profiles, and lower cardiovascular risk. Avocado, with its unique matrix of monounsaturated fats, antioxidants, and micronutrients, may represent a particularly potent functional food. Larger animal cohorts and mechanistic studies dissecting signaling pathways will be required to confirm these effects and guide translational trials. Nonetheless, our data provide compelling evidence that avocado supplementation can counteract L-NAME-induced hypertension and multi-organ injury, offering a promising nutraceutical adjunct for cardiovascular prevention and therapy.

## Conclusion

This study provides compelling preclinical evidence that avocado pulp supplementation offers multi-organ protection in a model of nitric oxide synthase inhibition. L-NAME administration induced significant hypertension, pathological coupling of systolic and diastolic pressures, hematological perturbations, hepatic stress, and renal dysfunction, consistent with the systemic impact of chronic vascular injury. Avocado supplementation, whether alone or in combination with pharmacological therapy, attenuated these abnormalities by normalizing blood pressure, disrupting maladaptive SBP–DBP correlations, reducing thrombocytosis, and preserving hepatic and renal biochemical indices. The protective profile extended across cardiovascular, hematological, hepatic, and renal domains, highlighting avocado’s pleiotropic mechanisms, likely mediated through antioxidant, anti-inflammatory, and vasodilatory pathways. These findings underscore avocado’s potential as a nutraceutical intervention capable of stabilizing vascular and organ function in the context of cardiovascular stress. While further studies with larger cohorts and mechanistic dissection are required, our results support translational exploration of avocado as an adjunct to conventional therapy for hypertension and related multi-organ complications.

## Future Directions

Although this study highlights the protective potential of avocado supplementation against L-NAME– induced cardiovascular and multi-organ injury, further work is required to advance translation. Larger animal cohorts are needed to validate the reproducibility of these findings and to clarify dose–response relationships. Mechanistic studies should dissect molecular pathways, particularly nitric oxide signaling, oxidative stress, and inflammatory cascades, that mediate avocado’s protective effects. Integrative omics approaches, including transcriptomics, metabolomics, and lipidomics, could uncover novel biomarkers of avocado action and systems-level reprogramming. Ultimately, carefully designed human intervention trials in populations at risk for hypertension or metabolic syndrome will be essential to determine clinical efficacy and safety, and to position avocado as a viable nutraceutical adjunct to standard therapies.

## Acknowledgements

We are grateful to the students of the Departments of Biochemistry and Human Nutrition for their support.

